# Understanding drivers of phylogenetic clustering and terminal branch lengths distribution in epidemics of *Mycobacterium tuberculosis*

**DOI:** 10.1101/2022.01.03.474767

**Authors:** Fabrizio Menardo

## Abstract

Detecting factors associated with transmission is important to understand disease epidemics, and to design effective public health measures. Clustering and terminal branch lengths (TBL) analyses are commonly applied to genomic data sets of *Mycobacterium tuberculosis* (MTB) to identify sub-populations with increased transmission. Here, I used a simulation-based approach to investigate what epidemiological processes influence the results of clustering and TBL analyses, and whether difference in transmission can be detected with these methods. I simulated MTB epidemics with different dynamics (latency, infectious period, transmission rate, basic reproductive number R_0_, sampling proportion, and molecular clock), and found that all these factors, except the length of the infectious period and R_0_, affect the results of clustering and TBL distributions. I show that standard interpretations of this type of analyses ignore two main caveats: 1) clustering results and TBL depend on many factors that have nothing to do with transmission, 2) clustering results and TBL do not tell anything about whether the epidemic is stable, growing, or shrinking. An important consequence is that the optimal SNP threshold for clustering depends on the epidemiological conditions, and that sub-populations with different epidemiological characteristics should not be analyzed with the same threshold. Finally, these results suggest that different clustering rates and TBL distributions, that are found consistently between different MTB lineages, are probably due to intrinsic bacterial factors, and do not indicate necessarily differences in transmission or evolutionary success.

## Introduction

In the last decade well beyond half a million bacterial genomes have been sequenced worldwide, about 7% of these from *Mycobacterium tuberculosis* (MTB) strains (Blackwell et al. 2021). One of the reasons behind these extensive sequencing efforts is the use of whole genome sequencing in molecular epidemiology. Molecular epidemiology studies of MTB (and of other pathogens) use microbial genome sequences sampled from different patients to investigate epidemiological dynamics such as transmission, relapses, and the acquisition and spread of antibiotic resistance (Hatherell et al. 2016, Guthrie & Gardy 2017, Nikolayevskyy et al. 2019). One of the most popular approaches to analyze this data is to cluster strains in groups based on their genetic distance. The identification of clustered MTB strains is commonly interpreted as evidence for recent local transmission, while patients infected with non-clustered strains (singletons) are thought to be novel introductions (i.e. patients that got infected somewhere else). Similarly, when studying the epidemiology of antibiotic resistance, clustered strains are considered as cases of transmission of resistance, while singletons are thought to be more likely to represent instances of resistance acquisition (Hatherell et al. 2016).

In MTB studies, clusters are often defined based on a single nucleotide polymorphism (SNP) threshold: strains with fewer SNPs than a given threshold are grouped together in the same cluster. However, this has been criticized, because it is not clear which value should be used (Stimson et al. 2018). A review of more than 30 publications concluded that a threshold of 6 SNPs could be used to identify cases of direct transmission (Nikolayevskyy et al. 2019), but other studies proposed different thresholds for different settings (see Table 1 in Hatherell et al. 2016 for some examples).

**Table 1.** Parameters and results for the different scenarios in the analysis of transmission dynamics (λ: transmission rate, *ε*: sampling rate, R_0_: λ/(*ε*+*σ)), σ*: death rate, *ψ:* rate of progression to infectiousness, π: molecular clock rate in expected nucleotide changes per site per year, 95% SNP threshold: the minimum SNP threshold for which at least 95% of samples are clustered in at least 95% of simulations, 100% SNP threshold: the minimum SNP threshold for which 100% of samples are clustered in at least 95% of simulations, 95% CI TBL: the confidence interval for the overall TBL distribution).

| Scenario (median infectious period (months) - $R_0$) | $\lambda$ | $\epsilon$ | $R_0$ | $\sigma$ | $\psi$ | $\pi$ | 95% SNP threshold | 100% SNP threshold | 95% CI TBL |
| --- | --- | --- | --- | --- | --- | --- | --- | --- | --- |
| Long infectious period, shrinking epidemic (17 - 0.9) | 0.45 | 0.5 | 0.9 | 0 | 1 | $8 \times 10^{-8}$ | 7 | 16 | 0-3 |
| Long infectious period, stable epidemic (17 - 1) | 0.5 | 0.5 | 1 | 0 | 1 | $8 \times 10^{-8}$ | 6 | 16 | 0-3 |
| Long infectious period, growing epidemic (17 - 1.1) | 0.55 | 0.5 | 1.1 | 0 | 1 | $8 \times 10^{-8}$ | 6 | 16 | 0-3 |
| Medium infectious period, shrinking epidemic (8 - 0.9) | 0.9 | 1 | 0.9 | 0 | 1 | $8 \times 10^{-8}$ | 5 | 16 | 0-3 |
| Medium infectious period, stable epidemic (8 - 1) | 1 | 1 | 1 | 0 | 1 | $8 \times 10^{-8}$ | 4 | 16 | 0-2 |
| Medium infectious period, growing epidemic (8 - 1.1) | 1.1 | 1 | 1.1 | 0 | 1 | $8 \times 10^{-8}$ | 4 | 14 | 0-2 |
| Short infectious period, shrinking epidemic (4 - 0.9) | 1.8 | 2 | 0.9 | 0 | 1 | $8 \times 10^{-8}$ | 5 | 14 | 0-2 |
| Short infectious period, stable epidemic (4 - 1) | 2 | 2 | 1 | 0 | 1 | $8 \times 10^{-8}$ | 4 | 15 | 0-2 |
| Short infectious period, growing epidemic (4 - 1.1) | 2.2 | 2 | 1.1 | 0 | 1 | $8 \times 10^{-8}$ | 3 | 16 | 0-2 |

Alternative approaches have been proposed to overcome the limits of SNP thresholds. For example, the method proposed by Stimson et al. (2018) groups strains based on a probabilistic model that takes into account the clock rate and the time of sampling. Other studies sidestepped the choice of one specific value by performing clustering multiple times with different thresholds, and comparing the results (Holt et. al. 2018, Meehan et al. 2018, Yang et al. 2018, López et al. 2020, Shuaib et al. 2020, Cox et al. 2021, Liu et al. 2021, Walter et al. 2021, Yang et al. 2021). Despite their limitations, clustering methods are considered useful to study transmission dynamics in MTB. Many studies tested the association of clustered strains with host and bacterial sub-populations, such as different age groups, HIV positive patients, bacterial lineages and others (Guerra-Assunção et al. 2015, Asare et al. 2020, Sobkowiak et al. 2020, Cox et al. 2021, Gygli et al. 2021, Merker et al. 2021, Yang et al. 2021). In these analyses, a positive association is interpreted as evidence for increased transmission of a certain sub-population. Further studies used clustering rates (the percentage of clustered strains), to characterize the extent of transmission in bacterial sub-populations (Holt et al. 2018, Shuaib et al. 2020). For example Holt et al. (2018) found that in Vietnam MTB Lineage 2 (L2) had higher clustering rates compared to MTB Lineage 4 (L4) and MTB Lineage 1 (L1), and interpreted these results as evidence for more frequent transmission of L2 strains, compared to L4 and L1 strains. Finally, in recent studies the distribution of terminal branch lengths (TBL) was used as a proxy for transmissibility in different MTB lineages (Holt et al. 2018, Freschi et al. 2021,Walter et al. 2021), partially complementing classical clustering analyses. All these approaches are based on the assumption that increased transmission results in increased clustering rates and shorter terminal branches. However, it is known that other factors could influence the results of clustering, for example it was posited that higher rates of molecular evolution, and low sampling rates should lead to lower clustering rates (Stimson et al. 2018, Menardo et al. 2019).

There is a consensus that epidemiological dynamics have an influence on the shape of MTB phylogenies, and therefore on clustering and TBL, though how they do so was never explored with quantitative studies. Here, I used simulations to explore what factors influence clustering results and TBL in MTB epidemics. I simulated the molecular evolution of MTB strains under different epidemiological conditions. I then inferred phylogenetic trees and computed clustering rates and TBL from the simulated data. With this approach I investigated the molecular clock rate, sampling proportion, transmission rate, basic reproductive number R_0_, length of the latency period, and length of the infectious period. I found that all these factors affect the results of clustering and TBL, with the exception of the lengths of the infectious period, and R_0_. These results are in contradiction with the standard interpretation of MTB epidemiological studies, namely that sub-populations associated with clustering and shorter terminal branches are necessarily transmitting more.

## Results

To investigate the expected patterns of genetic diversity under different epidemiological conditions, I assembled a pipeline to simulate the evolution of MTB genomes in different epidemiological settings (Fig. 1). The details are reported in the Methods section. Briefly, the pipeline simulates a transmission tree using an epidemiological model in which transmission events occur at rate λ and originate in the infectious compartment I, leading to a new exposed individual in compartment E. Exposed individuals become infectious at rate *ψ*, by moving from compartment E to compartment I. The time spent in the compartment E represents the latency period. Individuals are removed from compartment I by death or self cure, occurring at rate *σ*, or sampling, occurring at rate *ε*. The time spent in compartment I represents the infectious period.

**Figure 1.**
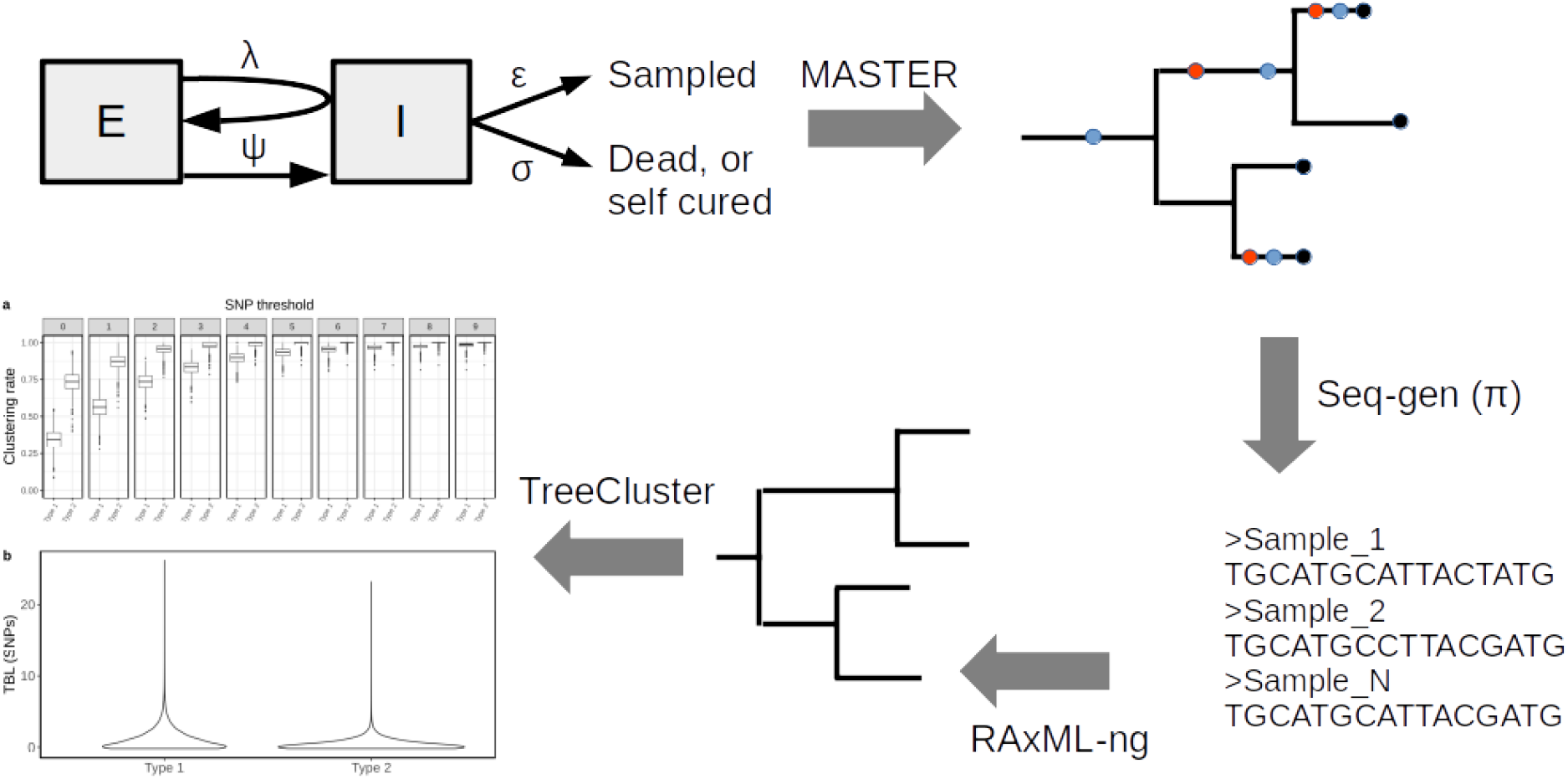
Simplified workflow, not all steps are depicted (see the Methods section for details). A transmission tree is simulated by MASTER, given the represented epidemiological model and a set of parameters (λ, *ψ, σ* and *ε))*. Seq-gen is used to simulate the evolution of MTB genome sequences along the tree, given a clock rate (π). RAxML-ng is used to estimate the phylogenetic tree from the sequence data, and TreeCluster to perform clustering.

A similar epidemiological model was used previously in a phylodynamic analysis of two MTB outbreaks (Kühnert et al. 2016, Kühnert et al. 2018). In a second step, the pipeline simulates the evolution of genome sequences along the tree given a clock rate π, and it computes the clustering rates under different SNP thresholds, and the the terminal branch lengths. The pipeline allows to test how clustering rates and TBL distribution change under different sampling proportions, molecular clock rates, transmission rates, basic reproductive numbers (R_0_= λ/(*σ*+*ε))*, lengths of the latency period (determined by *ψ*), and lengths of the infectious period (determined by *σ*+*ε*). Moreover, with this approach it is possible to investigate how the different factors impact the choice of a sensible SNP threshold. To do this, I defined the “95% sensitivity SNP threshold” as the minimum threshold for which at least 95% of strains are clustered in at least 95% of the simulations. A lower SNP threshold would lead to lower sensitivity (i.e. simulated samples which by definition are the result of recent transmission would not be clustered), while a larger threshold would lead to low specificity, although this cannot be quantified with this analysis.

### Factors influencing the results of clustering and TBL

First, I investigated whether epidemics with different basic reproductive numbers (R_0_) have different clustering rates and TBL, and whether strains from expanding populations are more likely to be clustered. In the epidemiological model used for this study, R_0_ corresponds to the ratio between the transmission rate (λ) and the sum of the death and sampling rates (*σ* and *ε*). For simplicity, in this analysis I set the death rate *σ* to zero, meaning that all cases are sampled. Therefore, the length of the infectious period is defined by the sampling rate. Typically the shift to infectiousness in TB patients is considered to occur with the onset of symptoms. However, in recent years the importance of sub-clinical TB has been reconsidered. It is possible that a considerable part of TB transmission occurs from asymptomatic patients, although this has not been yet quantified (Kendall et al. 2021). Given the uncertainty in the lengths of the infectious period I tested three different sampling rates: *ε =* 0.5, 1, and 2, corresponding to a median infectious period of ∼ 16.6, 8.3, and 4.2 months, respectively. These values cover well the possible length of the symptomatic period estimated in different countries (Ku et al. 2021). For each value of *ε*, I considered three different transmission rates leading to three scenarios: a shrinking epidemic with R_0_ = 0.9, a stable epidemic with R_0_ = 1, and a growing epidemic with R_0_ = 1.1. All other parameters were constant in all simulations. The exact settings for each simulated scenario are reported in Table 1. I performed 1000 simulations for each of the above scenarios and found no correlation between R_0_ and clustering rates, nor between R_0_ and TBL distributions (Fig. 2, Table 1). Clustering rates increased with larger transmission and sampling rates, while terminal branches tended to be shorter. However, non of these two measures was predictive of R_0_, and therefore cannot be used to determine whether the epidemic is growing or not (more on this below). To disentangle the effect of transmission and sampling rate I performed an additional set of simulations in which I explored different values of R_0_, changing alternatively the transmission rate or the sampling rate (Sup. Information). This analysis revealed that clustering results and TBL depend on the transmission rate, with higher values leading to higher clustering rates and shorter TBL. Conversely, the sampling rate had no major effect (Sup. Figs. 1 and 2, Sup. Table 1). To summarize the results so far, high transmission rates (per unit of time) result in larger clustering rates and shorter TBL. However these are not informative on whether an epidemic is growing or not, because this depends on the ratio between the transmission rate and the rate of removal from compartment I (*σ*+*ε*).

**Figure 2.**
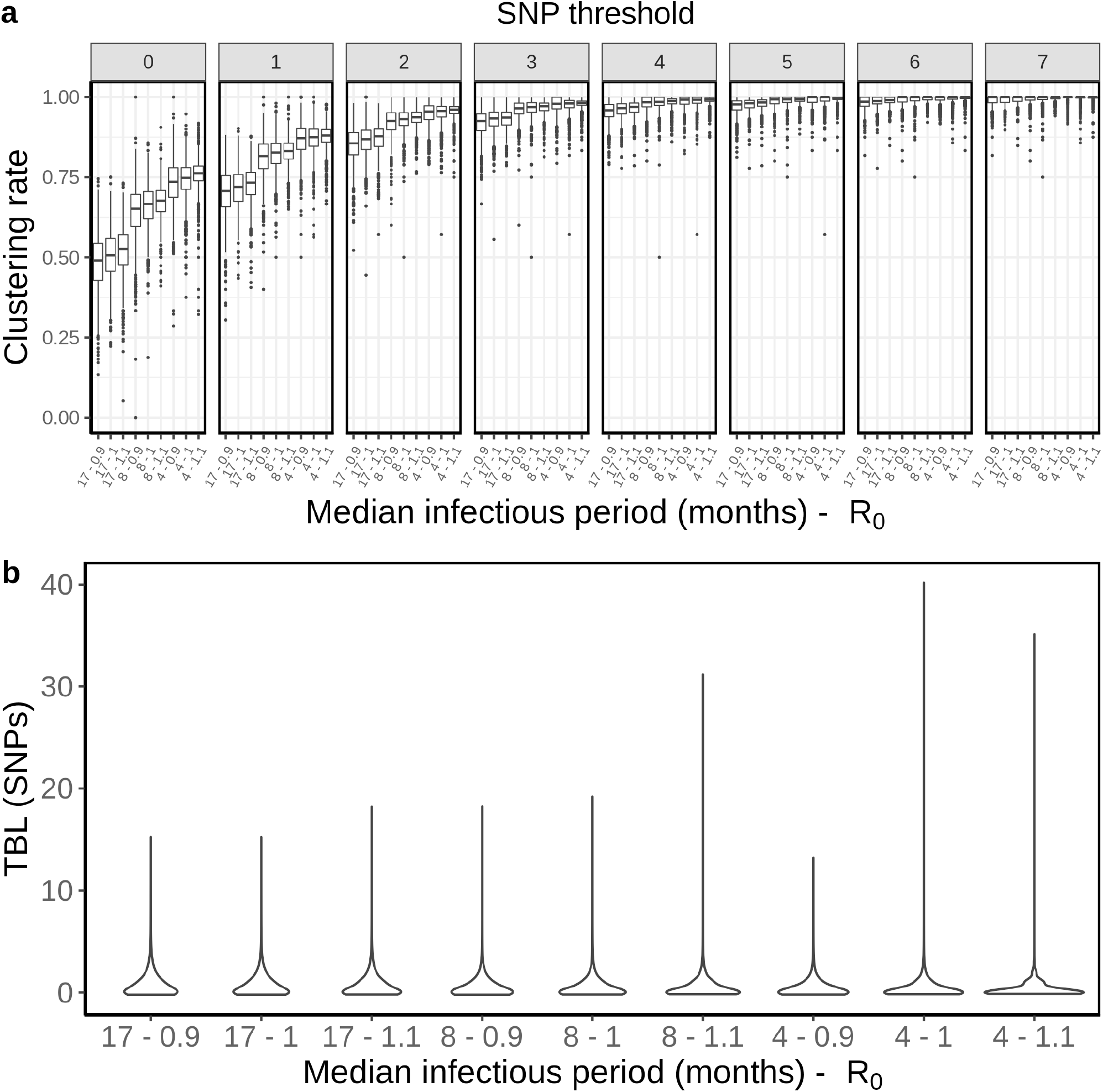
Clustering rates and TBL distributions for different transmission and sampling rates, resulting in different infectious periods and R_0_. **a)** Clustering rates with different SNP thresholds. Only SNP thresholds up to the highest 95% sensitivity threshold are plotted (i.e. for thresholds higher than 7 SNPs more than 95% of samples are clustered in more than 95% of simulations for all settings). **b)** Overall TBL distributions computed by merging all simulations.

Next, I used the same approach to explore other epidemiological and biological factors. I investigated different latency periods (defined by the rate of progression to infectiousness *ψ*), different rates of molecular evolution (π), and different sampling proportions (*ε*/(*σ*+*ε*)). The details of these analyses can be found as Supplementary Information. In a nutshell I found that: 1) longer latency periods resulted in lower clustering rates and longer TBL (Sup. Fig. 3, Sup. Table 2), 2) higher clock rates led to lower clustering rates and longer TBL (Sup. Fig. 4, Sup. Table 3), and 3) lower sampling proportion resulted in lower clustering rates and longer TBL (Sup. Fig. 5, Sup. Table 4). Finally, I also tested whether different sample sizes could have an influence on the results, and found that TBL and median clustering rates did not change when using a lower minimum number of tips in the simulated tree (Sup. Fig. 6, Sup. Table 5).

### An example

The results reported above have important implications on the interpretation of molecular epidemiology studies of MTB. To illustrate this in practice I will make an example in which two different strains of MTB are causing an epidemic in the same area. The two strains have different epidemiological and biological characteristics. We can think about it like two lineages of MTB, but it could also be two different clones belonging to the same lineage. MTB type 1 is expanding, with a R_0_ of 1.1, and it is characterized by a slow disease progression, with a median latency and infectious periods of about one year each. Type 1 populations are expected to increase by ∼95% every ten years. Conversely type 2 is shrinking, with a R_0_ of 0.9, and it is characterized by shorter median latency and infectious periods, approximately 5 months each. Type 2 populations are expected to shrink by ∼85% every 10 years. In addition, type 1 and 2 have moderate differences in their rate of molecular evolution, with a clock rate of respectively 1×10^−7^ and 7×10^−8^ nucleotide changes per site per year. Obviously, under this scenario type 1 is much more concerning for public health compared to type 2, at least on the long term. I repeated the same analysis presented above for the two types. The exact parameters for these simulations are reported in Table 2.

**Table 2.** Parameters and results for the two simulated scenarios in the practical example (λ: transmission rate, *ε*: sampling rate, R_0_ = λ/(*ε*+*σ)), σ*: death rate, *ψ:* rate of progression to infectiousness, π: molecular clock rate in expected nucleotide changes per site per year, 95% SNP threshold: the minimum SNP threshold for which at least 95% of samples are clustered in at least 95% of simulations, 100% SNP threshold: the minimum SNP threshold for which 100% of samples are clustered in at least 95% of simulations, 95% CI TBL: the confidence interval for the overall TBL distribution).

| Scenario | $\lambda$ | $\epsilon$ | $R_0$ | $\sigma$ | $\psi$ | $\pi$ | 95% SNP threshold | 100% SNP threshold | 95% CI TBL |
| --- | --- | --- | --- | --- | --- | --- | --- | --- | --- |
| Type 1 | 0.77 | 0.35 | 1.1 | 0.35 | 0.7 | $1 \times 10^{-7}$ | 9 | 21 | 0-5 |
| Type 2 | 1.53 | 0.85 | 0.9 | 0.85 | 1.7 | $7 \times 10^{-8}$ | 5 | 13 | 0-2 |

As expected, type 2 showed shorter terminal branches: the 95% CI of the TBL distribution was 0-5 and 0-2 SNPs for type 1 and type 2 respectively. Moreover, type 2 had higher clustering rates (Fig. 2), and the 95% sensitivity threshold was 9 and 5 SNPs for type 1 and 2 respectively. A typical interpretation of this data would be that type 2 is transmitting more than type 1, because of the shorter terminal branches, and the higher clustering rates. Alternatively, if we picked a specific threshold we would find that type 2 strains are associated with clustering (for lower thresholds), or that there are no difference in clustering among the two types (for higher thresholds). In any case a classic molecular epidemiology analysis would conclude that type 2 is transmitting more than type 1, or that there are no difference in transmission between the two types. While it is true that the transmission rate per unit of time is twice as high for type 2, it is also true that type 2 is bound to extinction, while type 1 is growing exponentially.

This example shows the pitfalls of TBL clustering analyses to study transmission in MTB epidemics. Admittedly, the simulation parameters for this example were picked to highlight the potential problems. However, the epidemiological characteristics of circulating MTB strains are normally not known, and the combination of parameters used here is entirely possible. The length of the infectious and latent periods and the clock rates are well within the range of values estimated with different data sets (Menardo et al. 2019, Ku et al. 2021). Overall, these results show that the standard interpretation of clustering results and TBL distributions can be badly misleading in some epidemiological settings.

**Figure 2.**
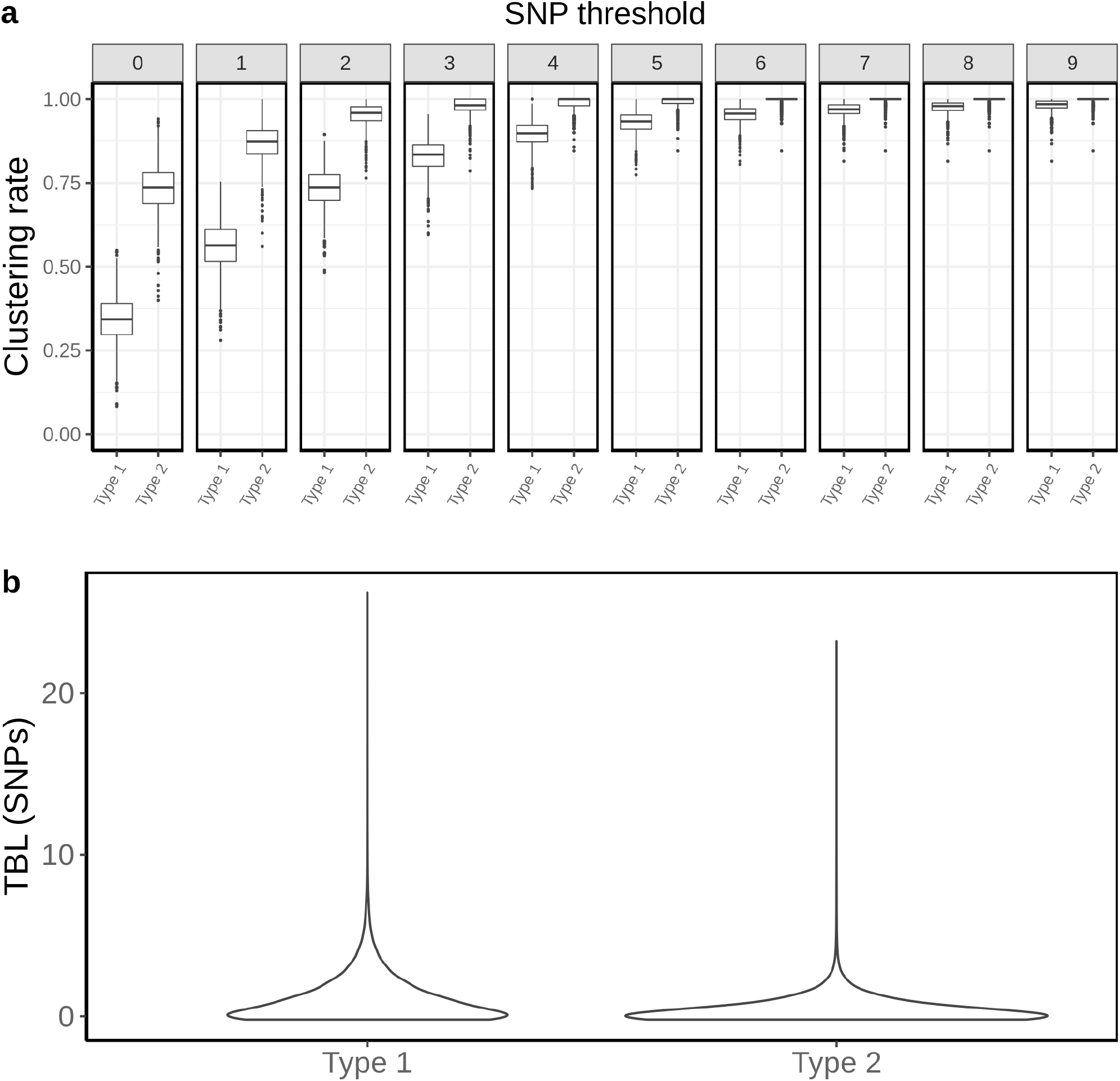
Clustering rates and TBL distributions for two different hypothetical sub-populations. Type 1 is expanding (R_0_ = 1.1), it has a long latency and infectious period (median: ∼12 months each), and a clock rate of 1×10^−7^. Type 2 has a R_0_ = 0.9, a short latency and infectious period (median: ∼5 months each), and a clock rate of 7×10^−8^. **a)** Clustering rates for the two types with different SNP thresholds. Only SNP thresholds up to the highest 95% sensitivity threshold are plotted (i.e. for higher thresholds more than 95% of samples are clustered in more than 95% of simulations for all settings). **b)** Overall TBL distributions computed by merging all simulations.

## Discussion

### The interpretation of clustering rates and TBL

Especially in low-incidence regions, clustering has proved to be useful to rapidly identify outbreaks and recent transmission (see Walker et al. 2018 for an example). However, the use of clustering analyses and their interpretation has evolved with time, and went beyond the identification of linked bacterial strains. Researchers often look for association with clusters, or differences in clustering rates, to characterize the extent of transmission in different sub-populations coexisting in high-incidence areas. Recently, the distribution of TBL has been used in a similar fashion. The results presented above demonstrate that these approaches suffer from two major limitations: 1) the transmission rate (per unit of time) does correlate positively with clustering rates, and negatively with TBL. However it is not the only factor doing so. The lengths of the latency period, the molecular clock rate and the sampling proportion all influence clustering and TBL, therefore difference between sub-populations might have nothing to do with difference in transmission. In other words, increased clustering might be due to shorter latency, lower clock rates or higher sampling proportion, and not to increased transmission. 2) The transmission rate expressed per unit of time does not determine the dynamic of an epidemic. Whether an epidemic, or a sub-population is growing or not, depends on the ratio between the transmission rate (λ) and the rate at which infectious individuals stop being infectious (*ε*+*σ*). This ratio (R_0_) represents the average number of transmission events during an individual infectious period (transmission per generation). R_0_ does not correlate with clustering rates or TBL, which can therefore not be used to determine whether a sub-population is stable, shrinking, or growing, neither in absolute, nor in relative terms (compared to a different sub-population). The confusion of transmission per unit of time and transmission per generation can lead to a mistaken interpretation of clustering and TBL analyses. These results resonate with the findings of Poon (2016), who reported that clustering methods used to study the epidemiology of HIV were biased towards detecting different sampling rates among sub-populations, and not variation in transmission rates.

### Is there an optimal SNP threshold?

The optimal SNP threshold is the one that maximize sensitivity and specificity. Here I defined the 95% sensitivity SNP threshold as the minimum threshold for which at least 95% of strains are clustered in at least 95% of the simulations. In other words, this is the threshold that maximize specificity at a 95% sensitivity level. One important result is that this threshold depends strongly on the epidemiological conditions: across all scenarios simulated for this study the 95% sensitivity threshold ranged between 3 and 11, and more extreme values are not impossible. Ideally, molecular epidemiological studies should use larger thresholds for regions characterized by longer latency, lower transmission rates, and/or lower sampling proportions. However, if the MTB population is not uniform, but consists of sub-populations with different epidemiological characteristics, or molecular clock rates, using a single SNP threshold will lead to biased results.

### Biology and epidemiology of MTB lineages

The issues discussed above are most relevant when comparing different bacterial sub-populations, such as the MTB lineages. Different MTB lineages have different clustering rates and distributions of TBL, independently from the region of sampling. For example, compared to other lineages, L2 was consistently found to have shorter TBL and higher clustering rates virtually everywhere, including in Vietnam (Holt et al. 2016, Le Hang et al. 2019), Malawi (Guerra-Assunção et al. 2015, Sobkowiak et al. 2020), Uzbekistan (Merker et al. 2018), South Africa (Cox et al. 2021), Georgia (Gygli et al. 2021), Iran (Vaziri et al. 2019), as well as globally (Freschi et al. 2021). At the opposite end of the spectrum, L1 was repeatedly found to have longer terminal branches, and lower clustering rates, compared to other lineages (Guerra-Assunção et al. 2015, Holt et al. 2016, Le Hang et al. 2019, Sobkowiak et al. 2020, Freschi et al. 2021). This repeated pattern is probably due to intrinsic bacterial factors, which affect clustering and TBL in all epidemiological settings. Different sampling proportions among lineages might play a role in some of these data sets, but it is unlikely that they are responsible for this widespread pattern. The three remaining factors that could explain the global differences between L2 and L1 are: 1) clock rates, 2) latency, and 3) transmission rates per unit of time.

1. A faster molecular clock for L1 would explain the lower clustering rates and longer terminal branches. However the little evidence that is available does not support this hypothesis. L1 and L2 were found to have similar clock rates when analyzed with the same set of methods, although the uncertainty of the estimates was very large (Menardo et al. 2019). Among other studies that inferred the clock rate for L2, most confirmed a moderately high rate (∼ 1×10^−7^, Merker et al. 2018, Rutaihwa et al. 2019, Torres Ortiz et al. 2021), while one estimated a higher rate of evolution (∼ 3×10^−7^, Eldholm et al. 2016). The only additional study to attempt the estimation of the clock rate of L1 found a lack of a temporal signal (Menardo et al. 2021). Altogether the available evidence is limited, and more precise estimates are needed to understand whether the rate of molecular evolution contributes to the different clustering rates and TBL for L1 and L2.
2. Longer latency would cause low clustering rates and long terminal branches in L1. Here, latency is defined as the period between being infected and becoming infectious, and it is typically thought to correspond to the period of asymptomatic infection. Although, pre-symptomatic transmission could be more important than previously thought (Kendall et al. 2021). There is some indirect evidence for a longer asymptomatic period in L1: a recent study estimated that asymptomatic infection is longer in TB patients in Southeast Asian countries such as Lao, Cambodia, and the Philippines (Ku et al. 2021), where L1 is responsible for more than half of TB cases (Schopfer et al. 2015, Chen et al. 2017, Somphavong 2018, Netikul et al. 2021). In these countries latency was found to be up to three times longer compared to Pakistan, Ethiopia and Zambia, where L1 is rare (Mulenga et al. 2010, Firdessa et al. 2013, Chihoota et al. 2018, Tulu & Ameni 2018, Wiens et al. 2018, Ali et al. 2019). Moreover, it is well documented that L2 is the most virulent MTB lineage (Hanekon et al. 2011, Peters et al. 2020), while L1 is less virulent compared to L2, L3 and L4 (Bottai et al. 2020). Virulence is often measured as bacterial growth rate in animal models or in human cell lines, and it seems quite natural that an infection of faster growing bacteria would have a shorter latency period.
3. A higher transmission rate per unit of time would also cause higher clustering rates and lower terminal branches in L2. This is how the results of these analyses are typically interpreted. Some partial evidence for this comes from a study in The Gambia, that found that household contacts of patients infected with L2 strains were more likely to develop disease within two years, compared to contacts of patients infected with other strains (De Jong et al. 2008). This could be the result of shorter latency, or higher transmission rates for L2. If L2 has a constitutively higher transmission rate, its proportion in a MTB population is expected to be stable only if the infectious period is shorter compared to other lineages. For example, in Malawi L2 was found to have higher clustering rates compared to L1, however the proportion of cases caused by the two lineages did not change over 20 years of monitoring (Sobkowiak et al. 2020). Assuming higher transmission rates for L2, these results can only be explained with a shorter infectious period of L2 compared to L1. Similarly to latency, the period between onset of symptoms and diagnosis was estimated to be longer in Southeast Asian countries where the MTB populations are dominated by L1 (Ku et al. 2021). This trend was not as strong as in the case of latency, but this is not surprising, as the delay in seeking care does not depend only on bacterial and host factors, but also on the public health system and other social conditions. As for latency, higher virulence could explain the higher transmission rate and shorter infectious period for L2 compared to L1.

To summarize, different clustering results and TBL between L1 and L2 are likely caused by difference in latency and or transmission rate per unit of time. However the analysis of TBL distribution and clusters can not tease these two factors apart. In this discussion I focused on L1 and L2, as these two lineages represent the extremes in term of clustering and TBL. For all other lineages the same logic applies, differences in latency, transmission and clock rates influence the tree topology in different combinations, resulting in intermediate TBL and clustering rates.

## Conclusions

The take home message of this study is that clustering analyses and TBL can tell us only so much about the epidemiological dynamics of MTB epidemics. While clustering will continue to be useful to detect linked strains, conclusions about differences in transmission among sub-populations are at best a simplification that conflate many different factors, at worst outright wrong. Phylodynamic methods that estimate the parameters of an epidemiological model from genomic data are becoming available (Kühnert et al. 2016, Didelot et al. 2017, Volz and Siveroni 2018). Although this type of analyses is challenging with current MTB data sets (Kühnert et al. 2018, Walter et al. 2021). Methodological advances, and more complete and longer sampling series of MTB epidemics will allow to study epidemiological dynamics more accurately in the future. In the meantime the results of clustering and TBL analyses should not be over-interpreted.

## Methods

MTB epidemics were simulated using an Exposed-Infectious epidemiological model. The model has two compartments, one for infectious individuals (I) and one for exposed individuals (E), the latter contains individuals that have been infected but are not yet infectious. Individuals in compartment I generate new infections adding new exposed individuals in compartment E with rate λ. Exposed individuals become infectious moving from compartment E to compartment I with rate *ψ*. Individuals are removed from compartment I either by death (or self cure), occurring at rate σ or sampling, occurring at rate *ε*. Under this model the reproductive number R_0_ corresponds to λ/(*σ*+*ε))*. Each of these events is modeled as a Poisson process, so that the probability of the event to occur through time is exponentially distributed. The mean and median of the exponential distribution are calculated respectively as 1/rate and ln(2)/rate. For example, with *ψ* = 1, the average and median latency periods are 1 and ∼0.69 years, respectively.

The BEAST v2.6.6 (Bouckaert et al. 2019) package MASTER v6.1.2 (Vaughan et al. 2013) was used to simulate phylogenetic trees under the parameters of the epidemiological model. Simulations were stopped after a 30 years maximum, or before, in case the simulated lineage went extinct. A post-simulation condition on the number of sampled tips in the tree was implemented by specifying the minimum and maximum number of sampled tips in the phylogenetic tree necessary to accept the simulation (unless differently specified in the text, the minimum number of tips was 100, the maximum was 2500). If the simulation was accepted, the tree was sub-sampled to the most recent 10 years of sampling, i.e. all tips sampled more than 10 years before the most recent tip were discarded. Additionally an outgroup was added to the phylogenetic tree. Seq-gen v1.3.4 (Rambaut & Grass 1997) was used to simulate genome sequences given the phylogenetic trees and a clock rate in expected nucleotide changes per site per year. One chromosome of 4 Mb was simulated for each tip, corresponding to the size of the MTB genome that is usually considered after discarding repetitive regions (Brites et al. 2018). Simulations were run under a HKY+Γ model with transition-transversion ratio k = 2 and shape parameter for the gamma distribution α = 1.

Variable sites were extracted from the whole genome alignments with SNP-sites v2.5.1 (Page et al. 2016), recording the number of invariant sites. Phylogenetic trees were inferred form the SNP alignment with RAxML-NG v0.9.0 (Kozlov et al. 2019) using a HKY model, and 2 starting trees (1 random, 1 parsimony). The maximum likelihood tree was rooted using the outgroup, and the branch lengths were converted in expected number of SNPs multiplying them by the alignment length (the number of SNPs). Finally, Treecluster v1.0.3 (Balaban et al. 2019) was used to obtain clusters under different SNP thresholds *t* = (0, 1 … 49, 50), so that the maximum pairwise distance between tips in the cluster is at most *t* (method “max”).

Unless differently stated, 1000 simulations were performed for all tested conditions. Clustering rates were computed for each simulation individually, while the TBL distribution were computed merging all simulations with the same set of parameters.

The pipeline is wrapped in a python script, using Biopython v1.78 (Cock et al. 2009) and ETE3 v3.1.2 (Huerta-Cepas et al. 2016) to handle sequences and trees. Plots were generated with the R package ggplot2 v3.1.1 (Wickham 2011).

The simulation pipeline, scripts and data to reproduce these results are available at https://github.com/fmenardo/sim_cluster_MTB.

## Aknowledgments

This work was founded by the SNF Ambizione grant PZ00P3_193473. Computation was performed on the ScienceCluster at the University of Zurich.

