## Supplementary Information (Sup. Text, Sup. Tables and Sup. Figures) for "Understanding drivers of phylogenetic clustering and terminal branch lengths distribution in epidemics of *Mycobacterium tuberculosis*"

Fabrizio Menardo

Department of Plant and Microbial Biology, University of Zurich. Switzerland.

**Transmission and sampling rates**

I tested how the transmission and sampling rates affect the results of clustering and the TBL distribution. First I used five different values for the transmission rate λ= 0.8, 0.9 ,1 , 1.1 ,1.2, keeping all other parameters constant ($\pi$ = 8x$10^{-8}$, $\sigma$ = 0, $\varepsilon$ = 1, $\psi$ = 1). Higher transmission rates resulted in larger clustering rates and shorter TBL (Sup. Fig. 1, Sup. Table1). The 95% sensitivity threshold ranged from 7 SNPs, for λ = 0.8, to 4 SNPs for λ= 1.2; while the 95% CI of the TBL distribution ranged from 0-3 to 0-2 for λ = 0.8 and 1.2 respectively. Next, I considered different lengths of the infectious period, to do this I used a constant transmission rate λ = 1, and five different values of the sampling rate (1.25,1.11111, 1, 0.90909, 0.83333) resulting in the same range of $R_0$ as in the previous analysis: $R_0$ = 0.8, 0.9, 1, 1.1 and 1.2. I found essentially no differences in the clustering rates for different lengths of the infectious period (Sup. Fig. 2). The 95% sensitivity threshold was equal to 6 SNPs for the two shorter infectious periods (i.e. the higher sampling rates), while it was 4 SNPs for the three longer infectious periods, although this was caused by a greater dispersion of clustering rates in the former two, and not to differences in the median values. Indeed the minimum threshold for which 100% of the samples were clustered in at least 95% of the simulations showed no particular trend (15, 16, 16, 14, and 15 SNPs from the shortest to the longest infectious period). Finally the 95% CI of the TBL distribution was 0-2 SNPs for all parameter values. These results show that different transmission rates resulted in different clustering rates and TBL, while the sampling rate had no major effect.

Supplementary Table 1. Parameters and results for the different simulated scenarios in the analysis of transmission and sampling rates (λ: transmission rate, *ε*: sampling rate, $R_0 = \lambda/(\varepsilon+\sigma)$, *σ*: death rate, *ψ:* rate of progression to infectiousness, π: molecular clock rate in expected nucleotide changes per site per year, 95% SNP threshold: the minimum SNP threshold for which at least 95% of samples are clustered in at least 95% of simulations, 100% SNP threshold: the minimum SNP threshold for which 100% of samples are clustered in at least 95% of simulations, 95% CI TBL: the confidence interval for the overall TBL distribution).

| Scenario | λ | ε | $R_0$ | σ | ψ | π | 95% SNP threshold | 100% SNP threshold | 95% CI TBL |
|---|---|---|---|---|---|---|---|---|---|
| Fixed *ε*, $R_0$=0.8 | 0.8 | 1 | 0.8 | 0 | 1 | $8x10^{-8}$ | 7 | 17 | 0-3 |
| Fixed *ε*, $R_0$=0.9 | 0.9 | 1 | 0.9 | 0 | 1 | $8x10^{-8}$ | 5 | 16 | 0-3 |
| Fixed *ε*, $R_0$=1.1 | 1.1 | 1 | 1.1 | 0 | 1 | $8x10^{-8}$ | 4 | 14 | 0-2 |
| Fixed *ε*, $R_0$=1.2 | 1.2 | 1 | 1.2 | 0 | 1 | $8x10^{-8}$ | 4 | 16 | 0-2 |
| $\varepsilon=\lambda_{IE}$, $R_0$=1 | 1 | 1 | 1 | 0 | 1 | $8x10^{-8}$ | 4 | 16 | 0-2 |
| Fixed $\lambda_{IE}$, $R_0$=0.8 | 1 | 1.25 | 0.8 | 0 | 1 | $8x10^{-8}$ | 6 | 15 | 0-2 |
| Fixed $\lambda_{IE}$, $R_0$=0.9 | 1 | 1.11111 | 0.9 | 0 | 1 | $8x10^{-8}$ | 6 | 16 | 0-2 |
| Fixed $\lambda_{IE}$, $R_0$=1.1 | 1 | 0.90909 | 1.1 | 0 | 1 | $8x10^{-8}$ | 4 | 14 | 0-2 |
| Fixed $\lambda_{IE}$, $R_0$=1.2 | 1 | 0.83333 | 1.2 | 0 | 1 | $8x10^{-8}$ | 4 | 15 | 0-2 |

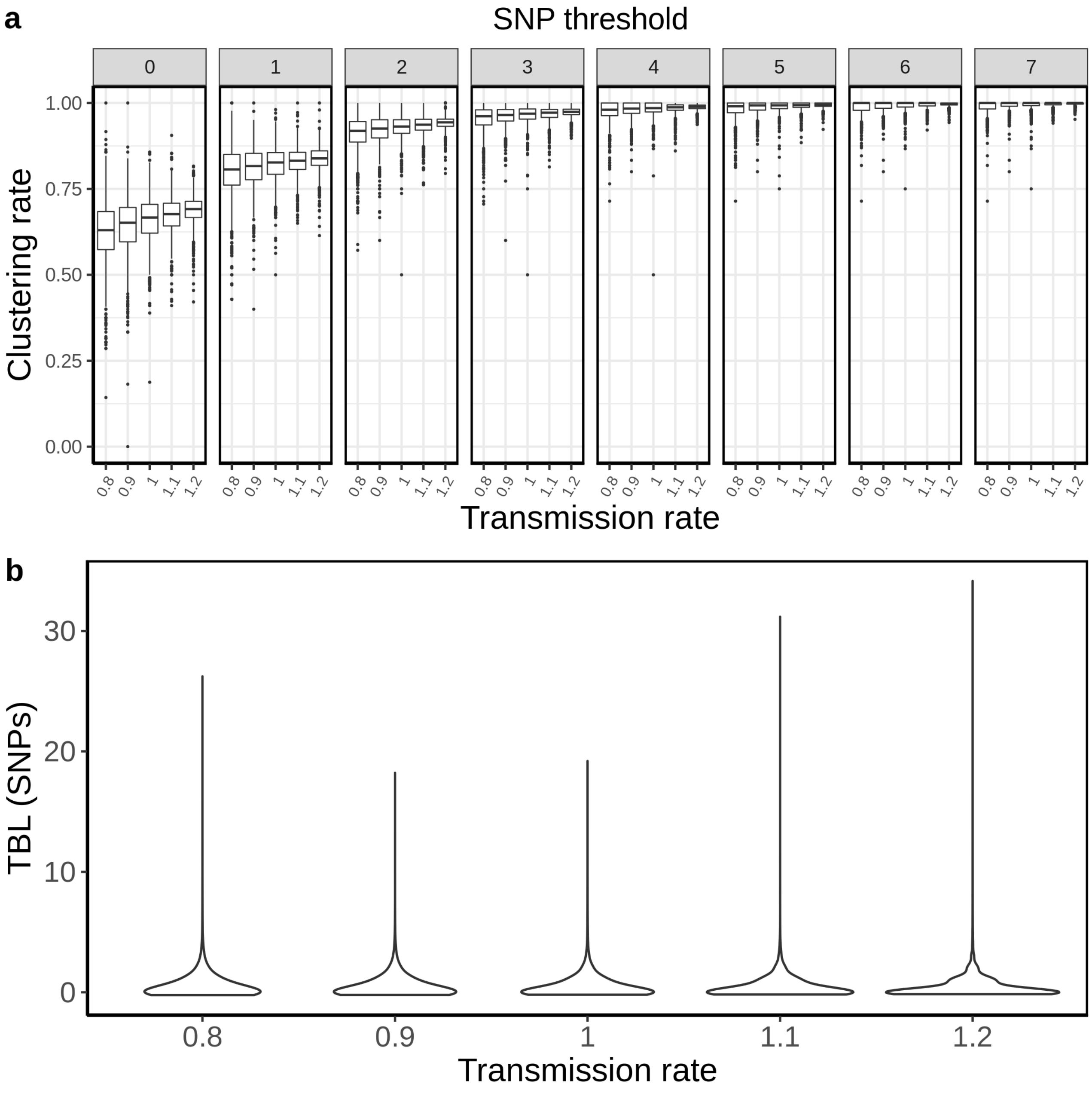


Supplementary Figure 1. Clustering rates and TBL distributions for different transmission rates. **a)** Clustering rates with different SNP thresholds. Only SNP thresholds up to the highest 95% sensitivity threshold are plotted (i.e. for higher thresholds more than 95% of samples are clustered in more than 95% of simulations for all settings). **b)** Overall TBL distributions computed by merging all simulations.

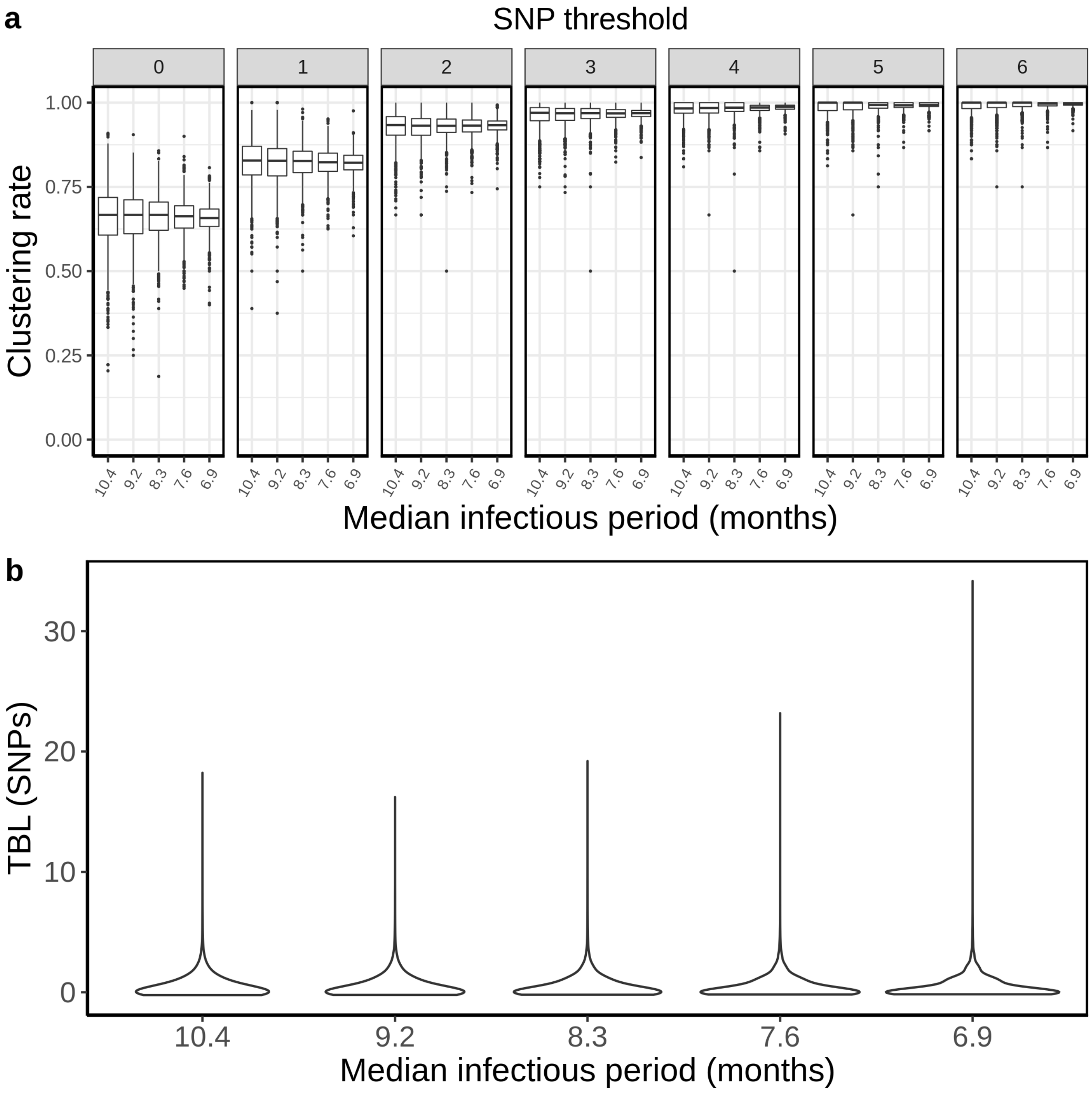


Supplementary Figure 2. Clustering rates and TBL distributions for different sampling rates, and therefore infectious periods. **a)** Clustering rates with different SNP thresholds. Only SNP thresholds up to the highest 95% sensitivity threshold are plotted (i.e. for higher thresholds more than 95% of samples are clustered in more than 95% of simulations for all settings). **b)** Overall TBL distributions computed by merging all simulations.

### Latency

I tested three different rates of progression to infectiousness $\psi$ = 0.5, 1, 2, corresponding to a median latent period of ~ 16.6, 8.3, and 4.2 months respectively. These values represent the range of duration of asymptomatic infection estimated in different countries (Ku et al. 2021). All other parameters were constant in all simulations ($\pi = 8x10^{-8}$; $\sigma = \varepsilon = 0.5$; $\lambda = 1$). Longer latency periods resulted in lower clustering rates and longer TBL (Sup. Fig 3). Correspondingly, the 95% sensitivity threshold were 10, 6, and 5 SNPs, and the 95% CI of the TBL distribution were 0-5, 0-3, and 0-2 SNPs, respectively for long, mid and short latency.

Supplementary Table 2. Parameters and results for the different simulated scenarios in the analysis of latency ($\lambda$: transmission rate, $\varepsilon$: sampling rate, $R_0 = \lambda/(\varepsilon+\sigma)$, $\sigma$: death rate, $\psi$: rate of progression to infectiousness, $\pi$: molecular clock rate in expected nucleotide changes per site per year, 95% SNP threshold: the minimum SNP threshold for which at least 95% of samples are clustered in at least 95% of simulations, 100% SNP threshold: the minimum SNP threshold for which 100% of samples are clustered in at least 95% of simulations, 95% CI TBL: the confidence interval for the overall TBL distribution).

| **Scenario** | **$\lambda$** | **$\varepsilon$** | **$R_0$** | **$\sigma$** | **$\psi$** | **$\pi$** | **95% SNP threshold** | **100% SNP threshold** | **95% CI TBL** |
|---|---|---|---|---|---|---|---|---|---|
| Short latency | 1 | 0.5 | 1 | 0.5 | 2 | $8x10^{-8}$ | 5 | 15 | 0-2 |
| Mid latency | 1 | 0.5 | 1 | 0.5 | 1 | $8x10^{-8}$ | 6 | 17 | 0-3 |
| Long latency | 1 | 0.5 | 1 | 0.5 | 0.5 | $8x10^{-8}$ | 10 | 20 | 0-5 |

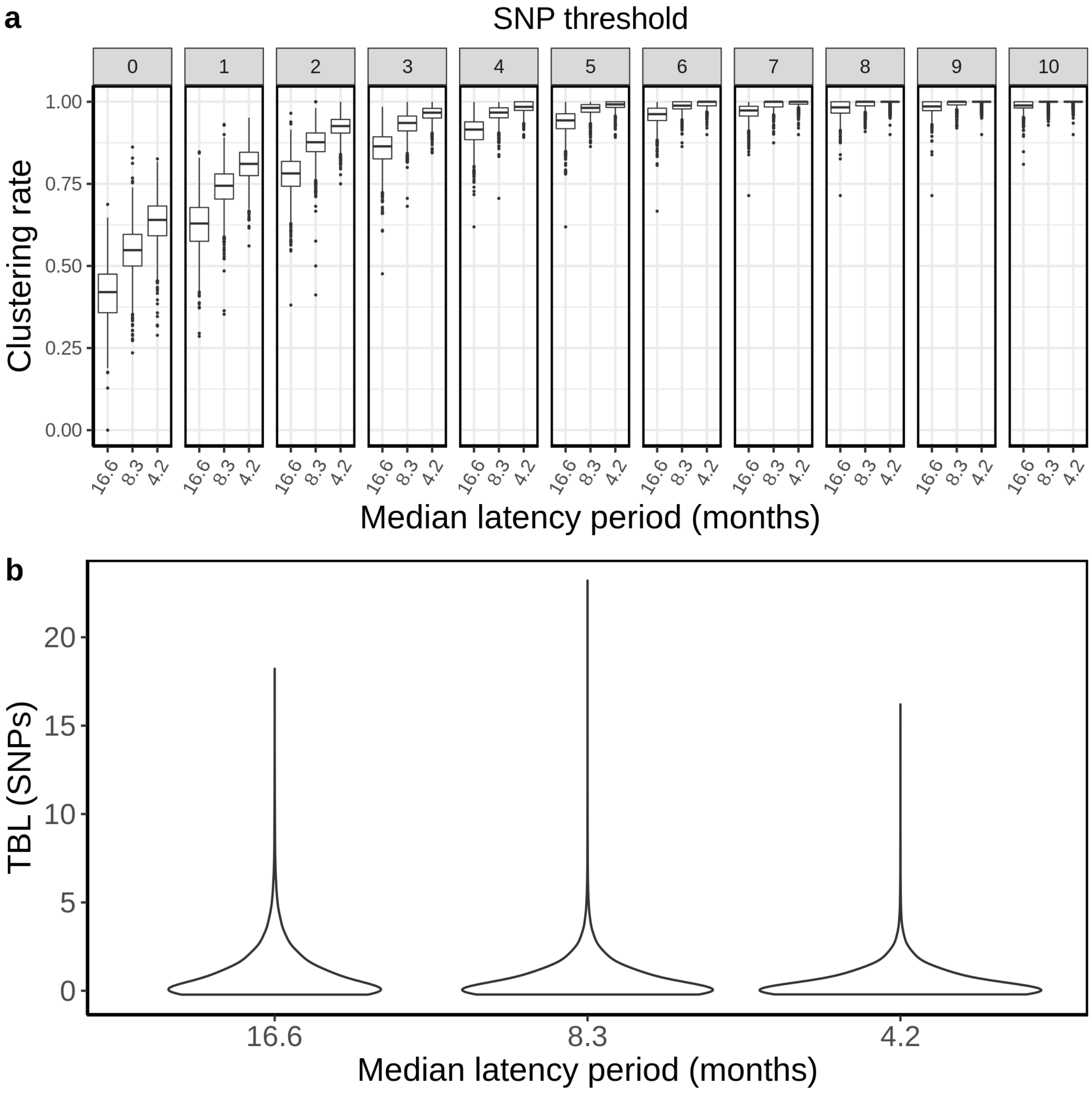


Supplementary Figure 3. Clustering rates and TBL distributions for different rates of progression to infectiousness, and therefore latency period periods. **a)** Clustering rates with different SNP thresholds. Only SNP thresholds up to the highest 95% sensitivity threshold are plotted (i.e. for higher thresholds more than 95% of samples are clustered in more than 95% of simulations for all settings). **b)** Overall TBL distributions computed by merging all simulations.

**Sampling proportion**

To test how different sampling proportions affect clustering rates and TBL, I used 4 different combinations of *σ* and *ε* (0.75 - 0.25; 0.5 - 0.5; 0.25 - 0.75; 0 - 1), corresponding to 4 sampling proportions: 25%, 50%, 75% and 100%. All other parameters were kept constant ($\pi = 8x10^{-8}$, $\lambda = \psi = 1$). Lower sampling proportions resulted in lower clustering rates and longer terminal branch lengths (Sup. Fig 4). The 95% sensitivity threshold was equal to 8, 6, 5, 4 for a sampling proportion of 25%, 50% 75% and 100% respectively, while the 95% CI of the TBL distribution were 0-4, 0-3, 0-3, and 0-2 (Sup. Table 3).

Supplementary Table 3. Parameters and results for the different simulated scenarios in the analysis of sampling proportions (λ: transmission rate, *ε*: sampling rate, $R_0 = \lambda/(\varepsilon+\sigma)$, *σ*: death rate, *ψ:* rate of progression to infectiousness, π: molecular clock rate in expected nucleotide changes per site per year, 95% SNP threshold: the minimum SNP threshold for which at least 95% of samples are clustered in at least 95% of simulations, 100% SNP threshold: the minimum SNP threshold for which 100% of samples are clustered in at least 95% of simulations, 95% CI TBL: the confidence interval for the overall TBL distribution).

| **Scenario** | **λ** | ***ε*** | **$R_0$** | ***σ*** | ***ψ*** | **π** | **95% SNP threshold** | **100% SNP threshold** | **95% CI TBL** |
|---|---|---|---|---|---|---|---|---|---|
| 25% sampling proportion | 1 | 0.25 | 1 | 0.75 | 1 | $8x10^{-8}$ | 8 | 15 | 0-4 |
| 50% sampling proportion | 1 | 0.5 | 1 | 0.5 | 1 | $8x10^{-8}$ | 6 | 17 | 0-3 |
| 75% sampling proportion | 1 | 0.75 | 1 | 0.25 | 1 | $8x10^{-8}$ | 5 | 20 | 0-3 |
| 100% sampling proportion | 1 | 1 | 1 | 0 | 1 | $8x10^{-8}$ | 4 | | 0-2 |

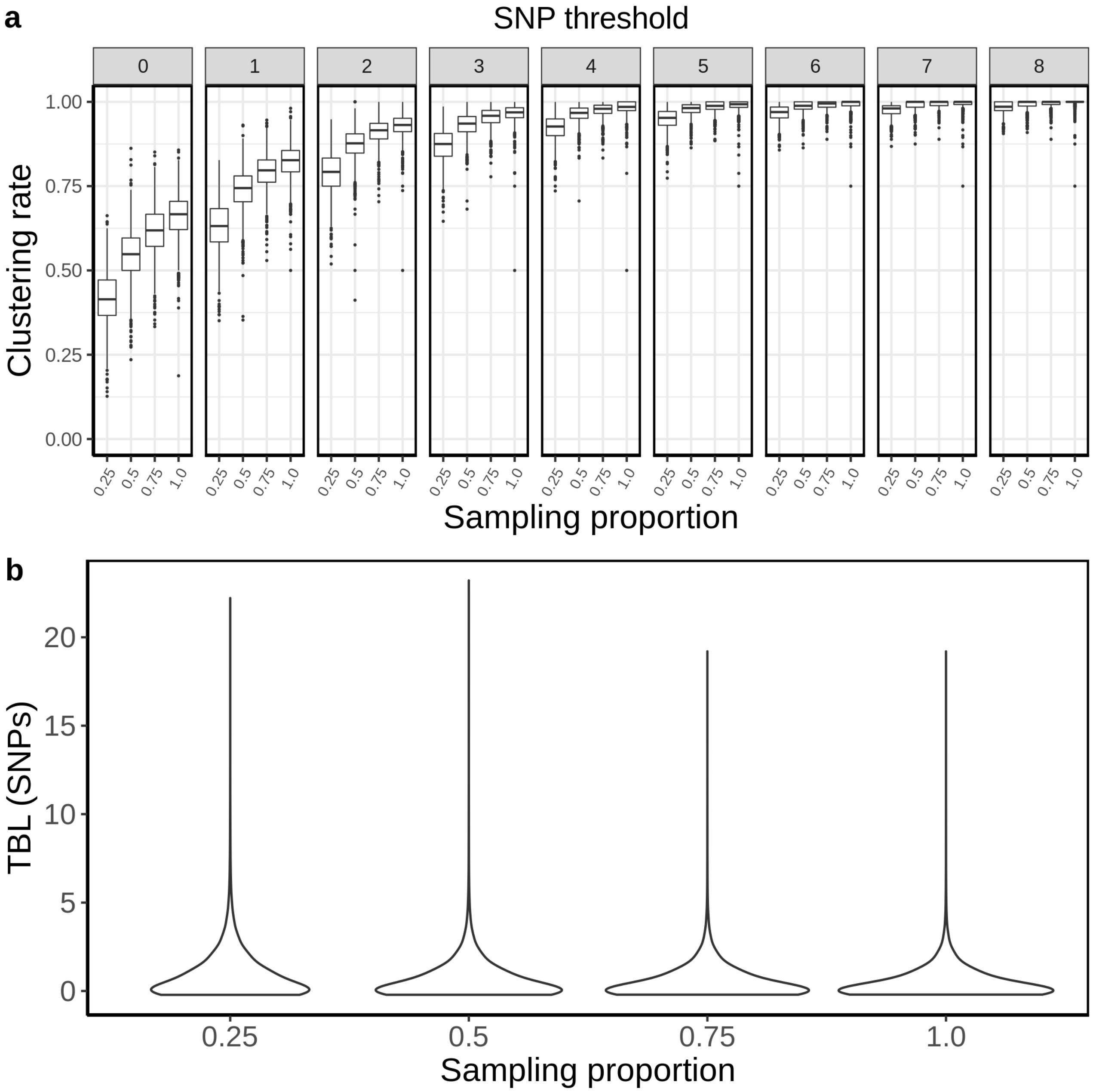


Supplementary Figure 4. Clustering rates and TBL distribution for different sampling proportions. **a)** Clustering rates with different SNP thresholds. Only SNP thresholds up to the highest 95% sensitivity threshold are plotted (i.e. for higher thresholds more than 95% of samples are clustered in more than 95% of simulations for all settings). **b)** Overall TBL distributions computed by merging all simulations.

**Molecular clock rates**

It is well known that faster evolving lineages will accumulated more mutations and therefore have longer TBL, and lower clustering rates (Stimson et al. 2019), although often this is overlooked. I tested three different rates of molecular evolution ($4x10^{-8}$, $8x10^{-8}$ and $1.2x10^{-7}$ nucleotide changes per site per year), roughly corresponding to the range of possible clock rates in MTB (Menardo et al. 2019). All other parameters were identical in all simulations ($\lambda = \psi = 1$; $\sigma = \varepsilon = 0.5$). For this analysis, I simulated 1000 transmission trees, and I then simulated the molecular evolution with different clock rates on the same set of trees. As expected, clustering rates were higher with low clock rates (Sup. Fig. 5), with the 95% sensitivity threshold equal to 4, 6 and 9, respectively for the lowest, mid, and highest clock rate. Also the TBL distributions were markedly different, with longer terminal branches for faster clock rates (95% CI = 0-2, 0-3 and 0-4, for low, mid and high clock rate).

Supplementary Table 4. Parameters and results for the different simulated scenarios in the analysis of clock rates ($\lambda$: transmission rate, $\varepsilon$: sampling rate, $R_0 = \lambda/(\varepsilon+\sigma)$, $\sigma$: death rate, $\psi$: rate of progression to infectiousness, $\pi$: molecular clock rate in expected nucleotide changes per site per year, 95% SNP threshold: the minimum SNP threshold for which at least 95% of samples are clustered in at least 95% of simulations, 100% SNP threshold: the minimum SNP threshold for which 100% of samples are clustered in at least 95% of simulations, 95% CI TBL: the confidence interval for the overall TBL distribution).

| **Scenario** | $\lambda$ | $\varepsilon$ | $R_0$ | $\sigma$ | $\psi$ | $\pi$ | **95% SNP threshold** | **100% SNP threshold** | **95% CI TBL** |
|---|---|---|---|---|---|---|---|---|---|
| Fast clock rate | 1 | 0.5 | 1 | 0.5 | 1 | $1.2x10^{-7}$ | 9 | 24 | 0-4 |
| Mid clock rate | 1 | 0.5 | 1 | 0.5 | 1 | $8x10^{-8}$ | 6 | 17 | 0-3 |
| Low clock rate | 1 | 0.5 | 1 | 0.5 | 1 | $4x10^{-8}$ | 4 | 9 | 0-2 |

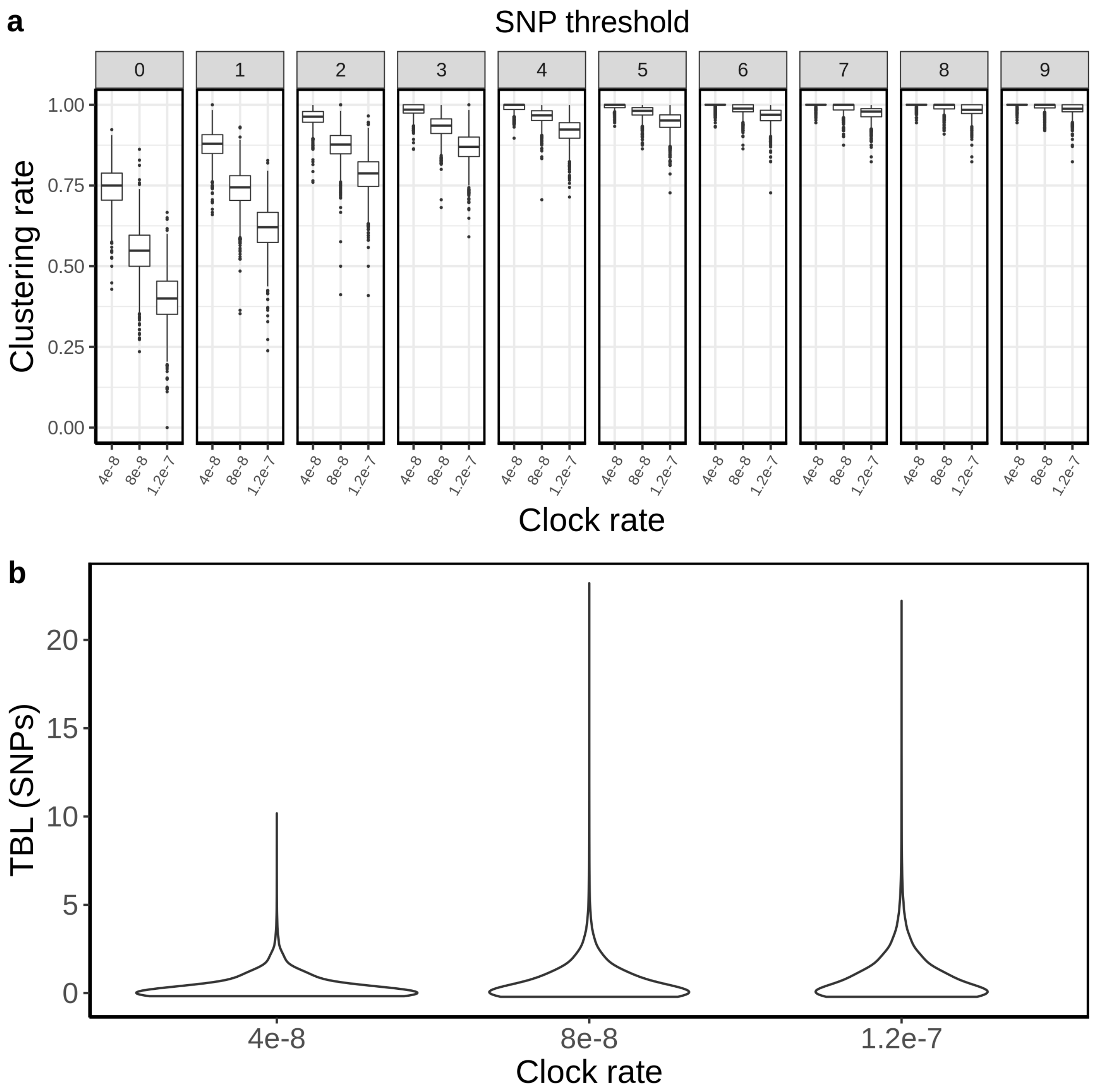


<u>Supplementary Figure 5.</u> Clustering rates and TBL distributions for different molecular clock rates. **a)** Clustering rates with different SNP thresholds. Only SNP thresholds up to the highest 95% sensitivity threshold are plotted (i.e. for higher thresholds more than 95% of samples are clustered in more than 95% of simulations for all settings). **b)** Overall TBL distributions computed by merging all simulations.

**Stop condition**

All results presented so far are based on the same simulation strategy: lineages are evolved for up to 30 years, or until they go extinct, and the last 10 years are sampled. The only condition imposed on a simulation to be accepted is a minimum and maximum number of tips in the transmission tree simulated by MASTER (before sampling the last 10 years). The maximum tip number was set to 2500, and in practice it is never reached within 30 years of simulated evolution, while the minimum number of tips was set to 100. I tested whether a smaller minimum number of tips could change the clustering rates and TBL. I used the same settings used in the mid clock rate analysis presented above, the only difference was that the minimum number of tips was set to 25, 50, or 100. The TBL distribution showed no difference (95% CI = 0-3 SNPs in all cases). The median of the clustering rates were also similar for all settings, the only difference was that the distribution of clustering rates was more dispersed with a lower number of tips. This caused the 95% sensitivity thresholds to be different: 11, 8, and 6 SNPs, respectively with 25, 50 and 100 minimum tips.
The larger dispersion was caused by the lower sample sizes, indeed the minimum threshold for which 100% of the samples were clustered in at least 95% of the simulations showed no particular trend (16, 17, and 17 SNPs from the lowest to the highest threshold).

Supplementary Table 5. Parameters and results for the different simulated scenarios in the analysis of the minimum number of tips to accept a simulation (λ: transmission rate, *ε*: sampling rate, $R_0 = \lambda/(\varepsilon+\sigma)$, *σ*: death rate, *ψ:* rate of progression to infectiousness, π: molecular clock rate in expected nucleotide changes per site per year, 95% SNP threshold: the minimum SNP threshold for which at least 95% of samples are clustered in at least 95% of simulations, 100% SNP threshold: the minimum SNP threshold for which 100% of samples are clustered in at least 95% of simulations, 95% CI TBL: the confidence interval for the overall TBL distribution).

| **Scenario** | **λ** | ***ε*** | **$R_0$** | ***σ*** | ***ψ*** | **π** | **95% SNP threshold** | **100% SNP threshold** | **95% CI TBL** |
|---|---|---|---|---|---|---|---|---|---|
| Min tips = 25 | 1 | 0.5 | 1 | 0.5 | 1 | $8x10^{-8}$ | 11 | 16 | 0-3 |
| Min tips = 50 | 1 | 0.5 | 1 | 0.5 | 1 | $8x10^{-8}$ | 8 | 17 | 0-3 |
| Min tips = 100 | 1 | 0.5 | 1 | 0.5 | 1 | $8x10^{-8}$ | 6 | 17 | 0-3 |

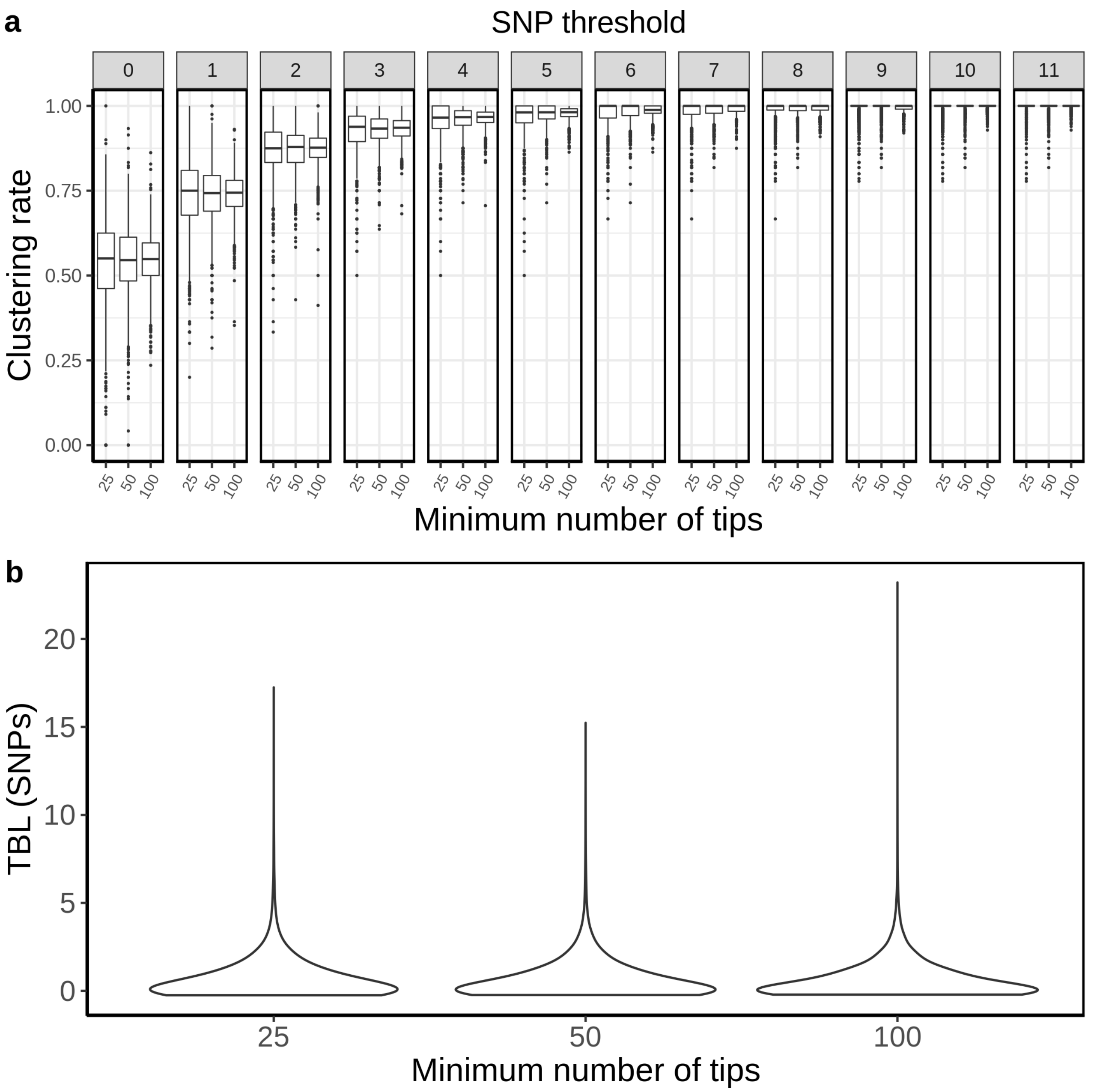


Supplementary Figure 6. Clustering rates and TBL distributions for different minimum number of tips necessary to accept the MASTER simulation. **a)** Clustering rates with different SNP thresholds. Only SNP thresholds up to the highest 95% sensitivity threshold are plotted (i.e. for higher thresholds more than 95% of samples are clustered in more than 95% of simulations for all settings). **b)** Overall TBL distributions computed by merging all simulations.

**References mentioned in Supplementary Information**

Ku, C. C., MacPherson, P., Khundi, M., Nzawa Soko, R. H., Feasey, H. R., Nliwasa, M., ... & Dodd, P. J. (2021). Durations of asymptomatic, symptomatic, and care-seeking phases of tuberculosis disease with a Bayesian analysis of prevalence survey and notification data. *BMC medicine*, *19*(1), 1-13.

Menardo, F., Duchêne, S., Brites, D., & Gagneux, S. (2019). The molecular clock of Mycobacterium tuberculosis. *PLoS pathogens, 15*(9), e1008067.

Stimson, J., Gardy, J., Mathema, B., Crudu, V., Cohen, T., & Colijn, C. (2019). Beyond the SNP threshold: identifying outbreak clusters using inferred transmissions. *Molecular biology and evolution*, *36*(3), 587-603.
